# Structure-based alignment of human caspase recruitment domains provides a framework for understanding their function

**DOI:** 10.1101/087908

**Authors:** Joseph P. Boyle, Tom Monie

## Abstract

Intracellular signalling is driven by protein-protein interactions. Members of the Death Domain superfamily mediate protein-protein interactions in both cell death and innate immune signalling pathways. They drive the formation of macromolecular complexes that act as a scaffold for protein recruitment and downstream signal transduction. Death Domain family members have low sequence identity, complicating their identification and predictions of their structure and function. We have taken all known human caspase recruitment domains (CARDs), a subfamily of the Death Domain superfamily, and generated a structure-guided sequence alignment. This alignment has enabled the identification of 14 positions that define the hydrophobic core and present a template for the identification of novel CARD sequences. We identify a conserved salt bridge in over half of all human CARDs and find a subset of CARDs likely to be regulated by tyrosine phosphorylation in their type I interface. Our alignment highlights that the CARDs of NLRC3 and NLRC5 are likely to be pseudodomains that have lost some of their original functionality. Together these studies demonstrate the benefits of structure-guided sequence alignments in understanding protein functionality.

## Introduction

The Death Domain protein superfamily contains a collection of helical protein domains that provide a central, and crucial, function in the formation of macro-molecular protein signalling complexes in both cell-death and immune signalling pathways. There are four members of the superfamily: the death domain (DD), the pyrin domain (PYD), the death effector domain (DED) and the caspase recruitment domain (CARD). Each of these protein domains folds into a helical bundle which is usually formed from six independent alpha helices (Kersse et al., 2011).

Most interactions between DD family members are homotypic in nature i.e. CARD with CARD. These interactions are mediated by discrete interfaces on the protein surface known as type I, II and III. Each of these interfaces consists of an ‘a’ and a ‘b’ component; one on each binding partner. For example, a Type I interaction involves the coming together of a Type Ia interface on one protein with a Type Ib interface on the other. The precise positioning of these interfaces means that potentially every DD-type fold involved in the complex can form six distinct interactions. It is this multiplicity of binding surfaces that helps to drive the formation of large signalling complexes such as the Myddosome (Lin et al., 2010; Motshwene et al., 2009) and Death Inducing Signalling Complex (Kischkel et al., 1995). Heterotypic interactions have been reported, but are not common. The first structure of a heterotypic DD interaction, between the CARD of RIPK2 (Receptor-Interacting Protein Kinase 2) and the DD of p75, has only recently been described (Lin et al., 2015). Interestingly, this complex was not formed by classical DD interfaces, suggesting that the heterotypic DD interactions may be somewhat more heterogeneous in nature.

Despite having very similar structures there is limited sequence similarity between, and even within, domain sub-families. Somewhat unusually it is not uncommon for two proteins in the same subfamily to have extremely similar tertiary structures, but possess less than 20% sequence identity. Consequently the identification of DD family members can be difficult, as can the reliable prediction of their secondary and tertiary structures.

Understanding how these protein domains interact is important for the potential modulation of signalling complexes involved in death and immune signalling networks. Amongst other functions the CARD plays an important role in the formation of the apoptosome (Cheng et al., 2016), recruitment of caspase-1 to the inflammasome (Guo et al., 2015), and propagation of signalling in the NOD1 (Bertin et al., 1999), NOD2 (Ogura et al., 2001), RIG-I and MDA-5 innate immune signalling pathways (Loo and Gale, 2011). In order to provide a framework by which we can better understand the functional diversity of the CARD we generated a structure-based alignment of all known human CARDs. This alignment identifies a broadly conserved hydrophobic core which functions as a signature motif for CARDs, provides a basis for the generation of homology models of unsolved CARD structures, and suggests the CARDs of human and murine NLRC3 and NLRC5 are pseudodomains.

## Results and Discussion

### Humans possess 36 CARD-containing and 4 partial/atypical CARD-containing proteins

To better understand the molecular basis of CARD:CARD mediated signalling pathways we sought to generate and analyse an alignment of human CARDs at the amino acid level. Sequence identification and retrieval began with the human PFAM collection (**PF00619**) which lists 121 human CARD-containing sequences, of which 29 are unique (Supplementary Table 1). By performing PSI-BLAST and jackhmmer searches with each of these unique sequences we expanded the collection of human CARDs to 36 through the addition of BINCA, CIITA, DLG5, MAVS, MDA-5 CARD1, NLRC3, RIG-I CARD1, RIG-I CARD2 and TNFRSF21 (Supplementary Table 1). None of PFAM, PSI-BLAST, or jackhmmer identified NLRC5 as a CARD-containing protein. However, the structure of the N-terminal domain of human NLRC5 has recently been solved (Gutte et al., 2014) and was classified as an atypical CARD. As such we added NLRC5 to our list of CARD-containing human proteins (Supplementary Table 1). Glutamine-Rich Protein 1 and NLRC3 were also labelled as partial CARDs because they do not appear to contain Helix 1. The TNFRSF21 sequence appears to be similarly distantly related to other members of the death domain superfamily and so was also treated as an irregular CARD. Consequently this resulted in the retrieval of 36 CARDs and 4 atypical human CARDs (Supplementary Table 1).

### Structure-guided alignment identifies a hydrophobic core of the CARD

Alignment of the CARD sequences was initially performed automatically using CLUSTALW2 (Larkin et al., 2007), CLUSTAL OMEGA (Sievers et al., 2011) and MUSCLE (Edgar, 2004). However, the outputs of these three programs were sufficiently different that we could not place any level of confidence in the accuracy of the alignment. In lieu of automatic alignment processes, we chose to use current structural information, available for 17 of the 36 typical CARDs (Supplementary Table 1), as a basis for the manual alignment of the human CARD sequences.

Initial comparative analysis of the available CARD structures indicated that the structures of Apaf-1, cIAP1, NLRP1, ASC and CARD8 could most easily be aligned due to the similar lengths of loops and insertions between their helices. The RAIDD CARD structure could be easily inserted into this alignment as its shorter helix 1 is complemented by an extended 1-2 loop (Figure 1A). As protein folding and stability is often driven by hydrophobic factors we determined which residues in our six aligned CARD structures composed the hydrophobic core of the domain. This identified 14 specific sites, all of which contained hydrophobic residues in each of the structures, and which we have labelled h1 to h14 (Figure 1B and C). Eleven of these residues are consistent with an earlier comparison of 11 different CARD structures, with residues h3, h8 and h13 representing an expansion to the defining hydrophobic of the CARD (Chen et al., 2013). Almost all of these conserved positions are positioned on the inside of amphipathic helices, however, residues h3 and h7 are part of the H1-H2 and H2-H3 loops respectively.

**Figure 1.**
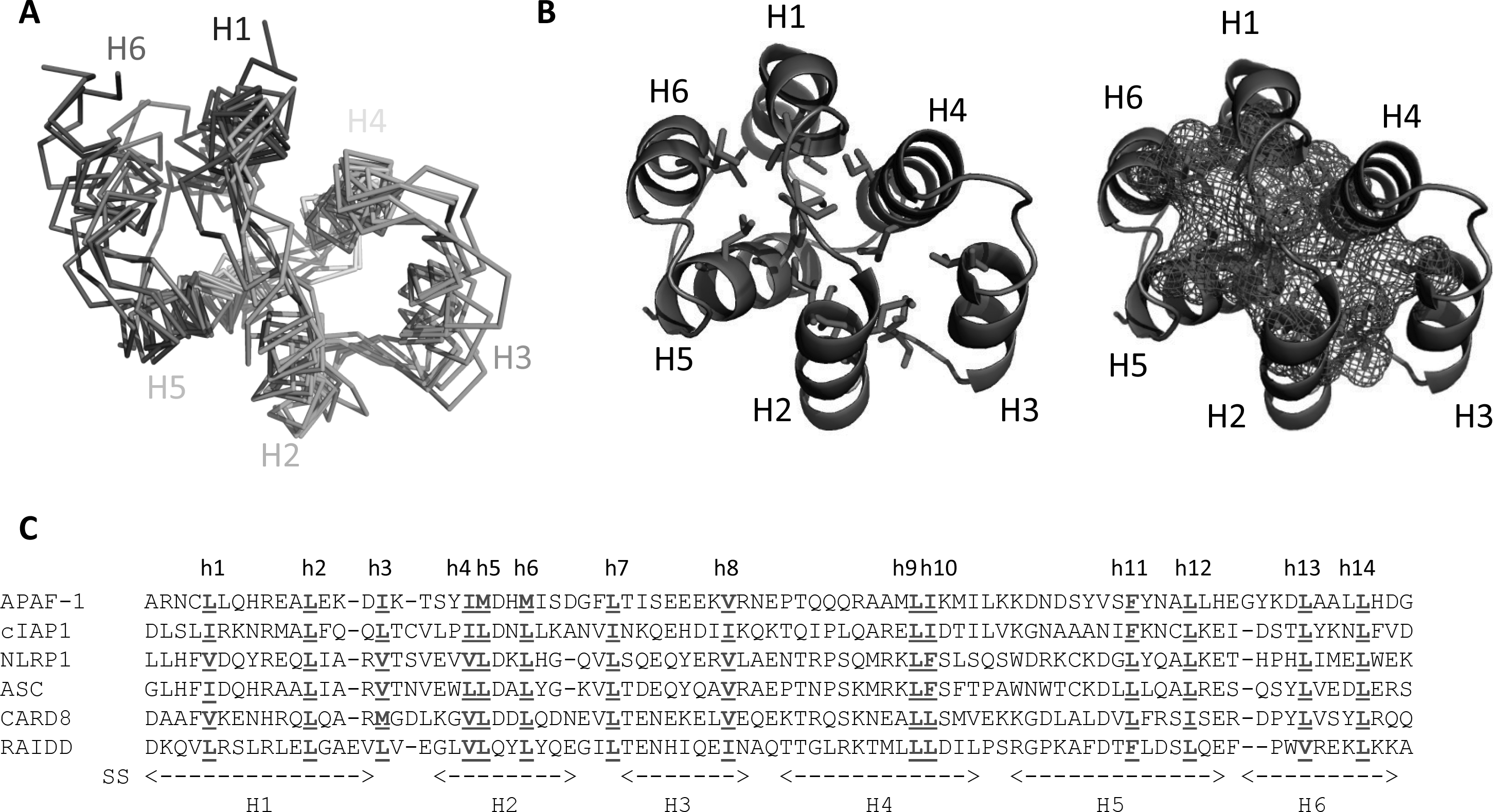
Alignment of CARD structures identifies a hydrophobic core. (**A**) Ribbon diagram of the superposition of the structures of Apaf-1 (PDB: 3YGS), cIAP1 (PDB: 2L9M), NLRP1 (PDB: 3KAT), ASC (PDB: 2KN6), CARD8 (PDB: 4IKM), and RAIDD (PDB: 3CRD). All sequences are coloured from blue at the N-terminus to red at the C-terminus. Individual helices are labelled and the greatest structural and spatial divergence is seen in helix 6. (**B**) Residues composing the hydrophobic core (coloured red, stick representation) of the CARD are plotted on the structure of the Apaf-I CARD (PDB: 3YGS). The right hand image uses a mesh to provide a spatial representation of the hydrophobic core. Individual helices are labelled. (**C**) Structure-based sequence alignment of six CARDs with highly similar overall structures. The positions of the residues contributing to the hydrophobic core are marked in red, bold and underlined, and labelled h1 to h14. The position of the six helices in the Apaf-I CARD is denoted below the sequence alignment.

### Generation of a complete structure-based alignment of the human CARDs

To generate a comprehensive structure-guided alignment of the human CARDs we sequentially compared the other known human CARD structures to this preliminary alignment. We began with the CARD of NOD1 which, consistent with other reports (Srimathi et al., 2008), immediately revealed a significant discrepancy between the first reported Nuclear Magnetic Resonance (NMR) structure of the NOD1 CARD (PDB 2B1W; (Manon et al., 2007)) and the subsequent crystal (PBDs: 4E9M, 2NZ7, 2NSN) and NMR structures (PDB: 2DBD) (Figure 2A and B). The NOD1 crystal structures exhibit a potentially non-physiological dimerisation involving helix swapping and so the most recent NMR structure of monomeric NOD1 (PDB: 2DBD) was used for alignment. NOD1 showed a two-residue insertion in the H5-H6 loop and a tilt in the orientation of helix 6 when compared to our six core structures (Figure 2C and D).

**Figure 2.**
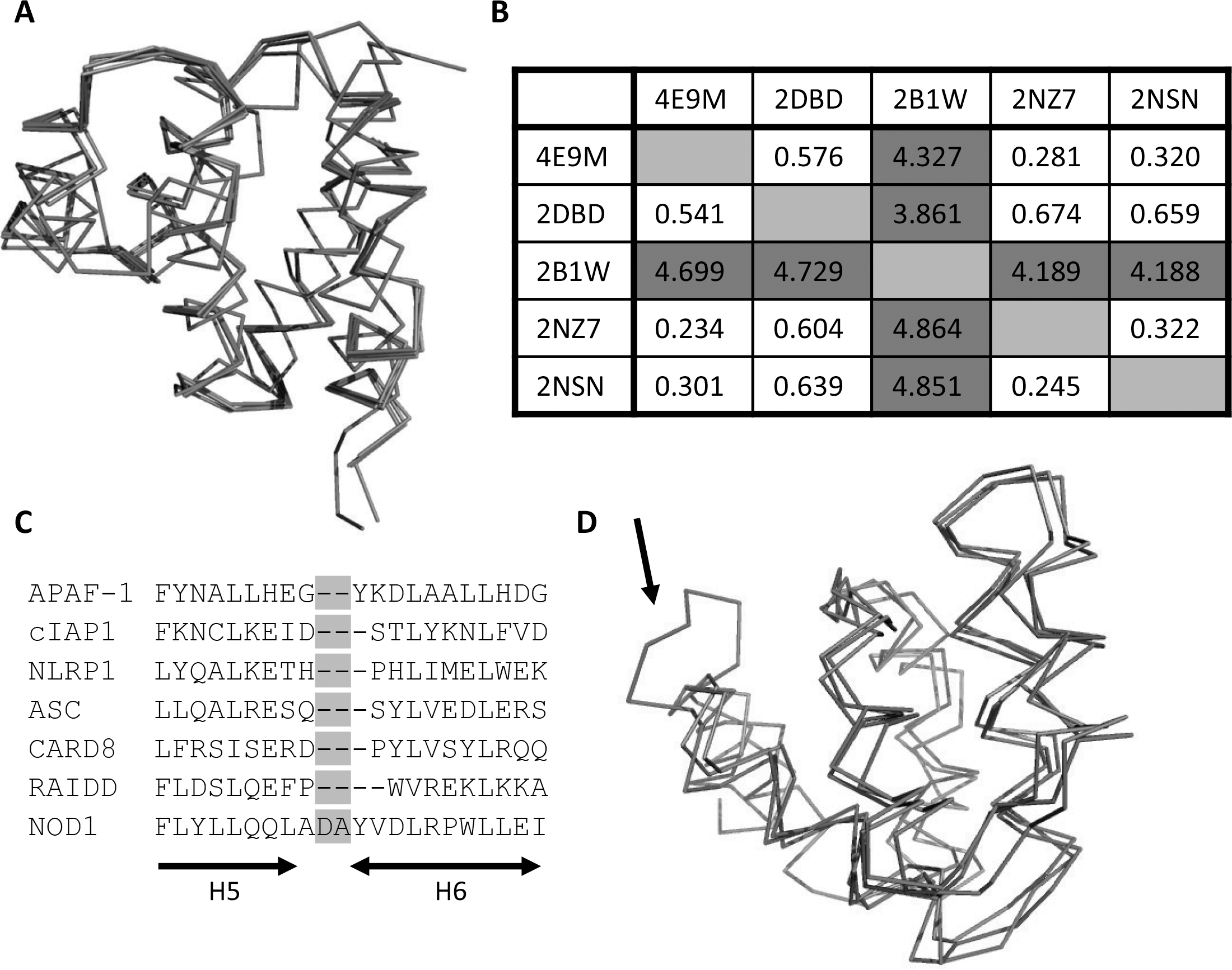
Comparison of Five NOD1 CARD Structures. (A) Superposition of the Cα backbone of helices 1-5 of the NOD1 CARD. The crystallographic NOD1 structures (PDBs: 4E9M, 2NZ7, 2NSN) and the NMR structure (PDB: 2DBD) are shown in grey. The original NMR structure (PDB: 2B1W) is shown in red. (**B**) Pairwise RMSD (Å) values for the superpositions of the NOD1 CARD structures shown in **A**. The upper triangle represents the RMSD of the full length CARDs, the lower triangle denotes the RMSD from just helices 2-5. The RMSD values for alignment against 2B1W are marked in red. (**C**) Alignment of the C-terminus of the NOD1 CARD with the initial six aligned structures. The insertion in the H5-H6 loop is highlighted in cyan. (**D**) Overlay of the NOD1 structure (red; PDB: 2DBD) with that of Apaf-1 (light grey; PDB: 3YGS) and cIAP1 (light grey; PDB: 2L9M) highlighting the structural impact of the extended H5-H6 loop (denoted by an arrow).

The assembly of our structure-based human CARD alignment was continued by the sequential manual addition of all the remaining available human CARD structures. This led to a number of minor modifications to the original alignment in order to incorporate the presence of additional insertions and deletions in the CARD sequences. CARD18 (also known as ICEBERG; PDB: 1DGN) for example, contains slightly extended H1-H2 and H3-H4 loops, whilst CARD11 shares a deletion in the H1-H2 loop with Apaf-1, but has an additional four residues in its H3-H4 loop (Figure 3). Caspase-9 (PDB: 4RHW) also possesses an insertion in the H3-H4 loop, but it is only two residues long (Figure 3). MAVS (PDB: 2VGQ) showed single residue deletions in two locations in the H1-H2 loop and also another in helix 5, as well as a three residue deletion at the end of helix 2 and a single residue insertion analogous to Apaf-1 in the final helix. It also had a three residue insertion in the H3-H4 loop. These elements were broadly similar to those seen in the two CARDs of RIG-I (PDB: 4P4H) except that in these instances the helix 2 deletion was only two residues long and the insertion in the H3-H4 loop for the first RIG-I CARD only contained two residues. The CARD of BINCA (PDB: 4DWN) showed four different insertions relative to the original alignment which served to extend the H1-H2 loop; extend the start and end of helix 2; and resulted in the breaking of helix 3 and the related introduction of an extended H3-H4 loop (Figure 4). Helix 6 was unstructured in RIPK2 (PDB: 2N7Z) and missing in NOL3 (PDB: 4UZ0). In fact, overall helix 6 showed variability between CARDs in its length, its position relative to helices 1-5, and its contribution to the hydrophobic core. This suggests that it is helices 1-5 that provide the critical structural and biological functionality of the CARD family.

**Figure 3.**
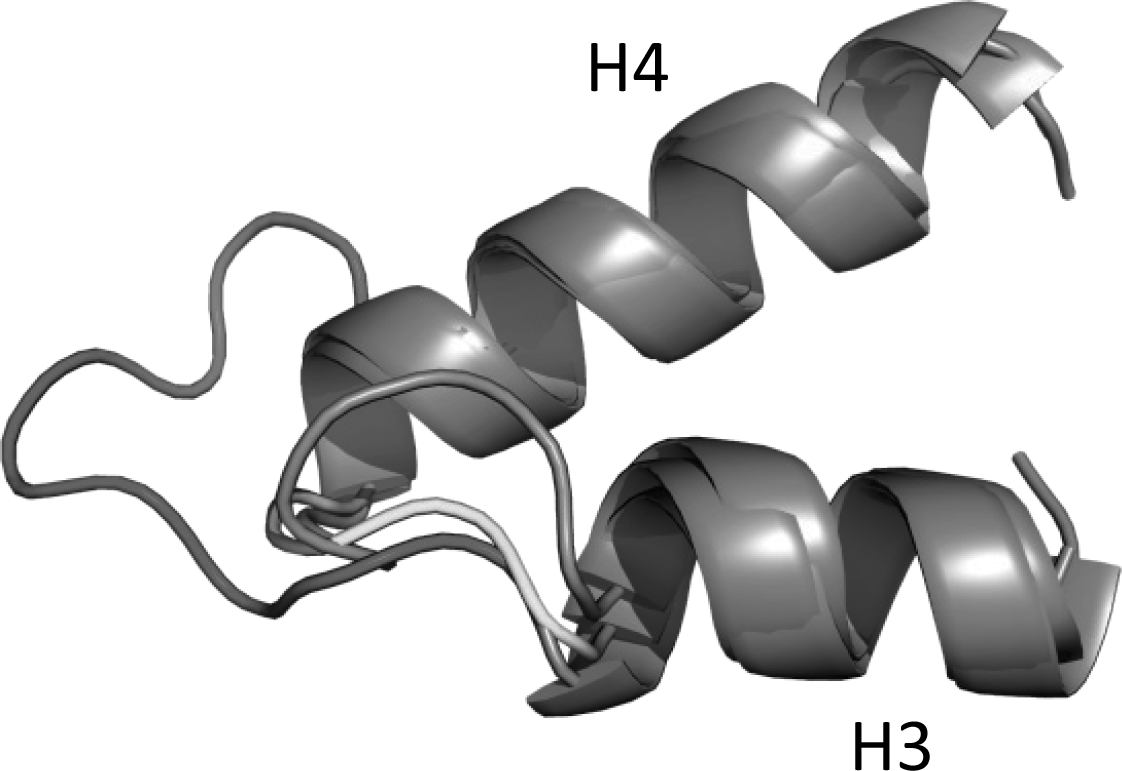
Extension of the H3-H4 loop in the Caspase-9 and CARD11 CARDs. Overlay of helix 3 (H3), the H3-H4 loop, and helix 4 (H4) from the CARDs of Apaf-1, caspase-9 and CARD11. The H3-H4 loops are coloured as follows: Apaf-I (yellow), caspase-9 (orange), and CARD11 (red).

**Figure 4.**
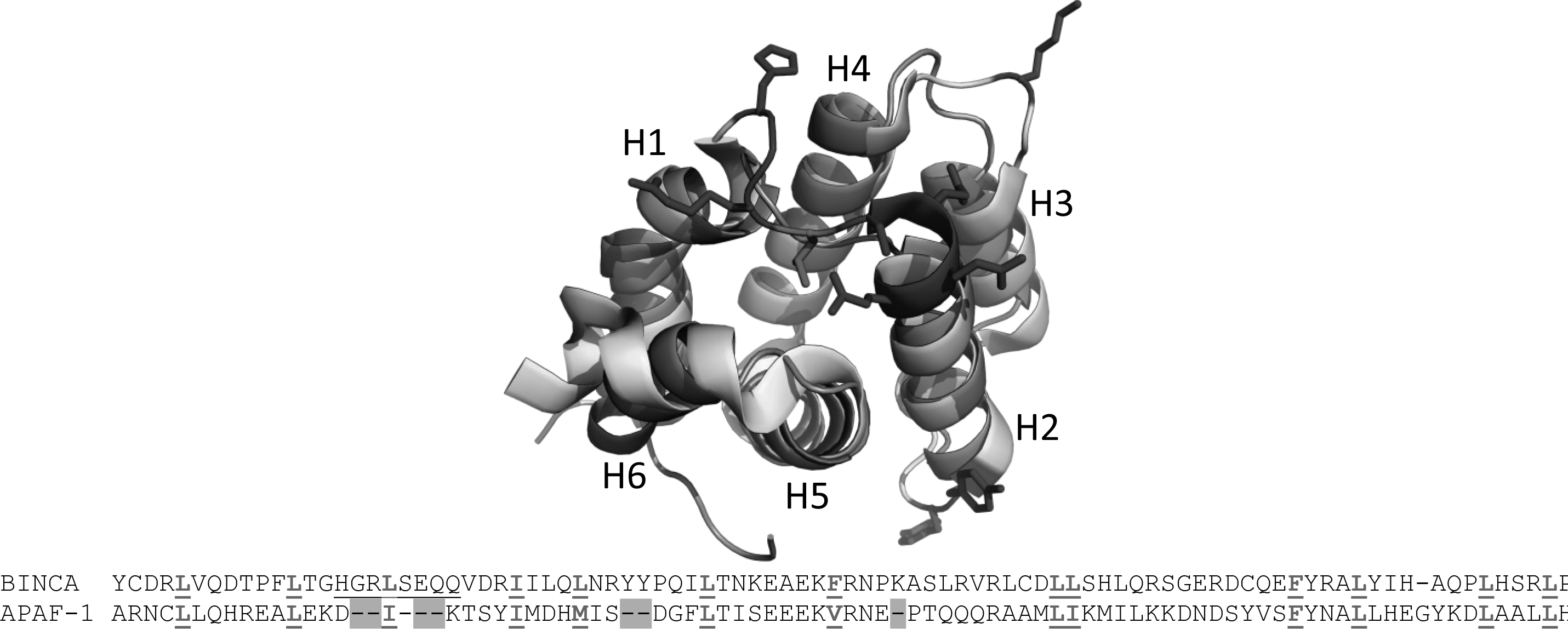
BINCARD contains insertions in the N-terminal half of its CARD. Superposition of the CARDs of Apaf-I (silver) and BINCARD (gold). Residues in BINCARD corresponding to the insertions in loops H1-H2, H2-H3 and H3-H4, as well as the extension of the N-terminus of Helix 2 are coloured blue and presented as sticks. The conserved hydrophobic position in the H1-H2 loop is coloured red. In the sequence alignment all core hydrophobic positions are coloured red and highlighted in bold and underlined. The gaps in Apaf-1 resulting from the BINCARD insertions are highlighted in cyan.

In total 17 different CARDs were used to generate a structure-guided alignment of the human CARD repertoire (Supplementary Figure 1). Aside from the absence of helix 6 in RIPK2 and NOL3 discussed above the hydrophobic core residues were conserved with the exception of residue h4. Despite pointing into the hydrophobic core this residue is a serine in the RIPK2 structure and a histidine in the Bcl-10 structure. This suggests that, at least in some positions, there is flexibility in the composition of these residues, though it remains to be determined whether there is any functional relevance to this variation. It should also be noted that the inherent structural variations in helix 6 means that these conserved hydrophobic residues could not always be assigned with complete confidence, but there tended to always be two hydrophobic residues facing towards the protein core.

### All typical human CARDs can be incorporated into the alignment

Once all the human CARDs with solved structures had been aligned we added in the sequences of the remaining 19 human CARDs using the presence of the core hydrophobic residues as a primary guide for the alignment, complemented using the FUGUE fold recognition server. The resulting alignment can be seen in Figure 5. NLRC3, NLRC5, GRP1 and TNFRSF21 were not included in the alignment due to their extensive deviation from the classical CARD sequence and structure.

**Figure 5.**
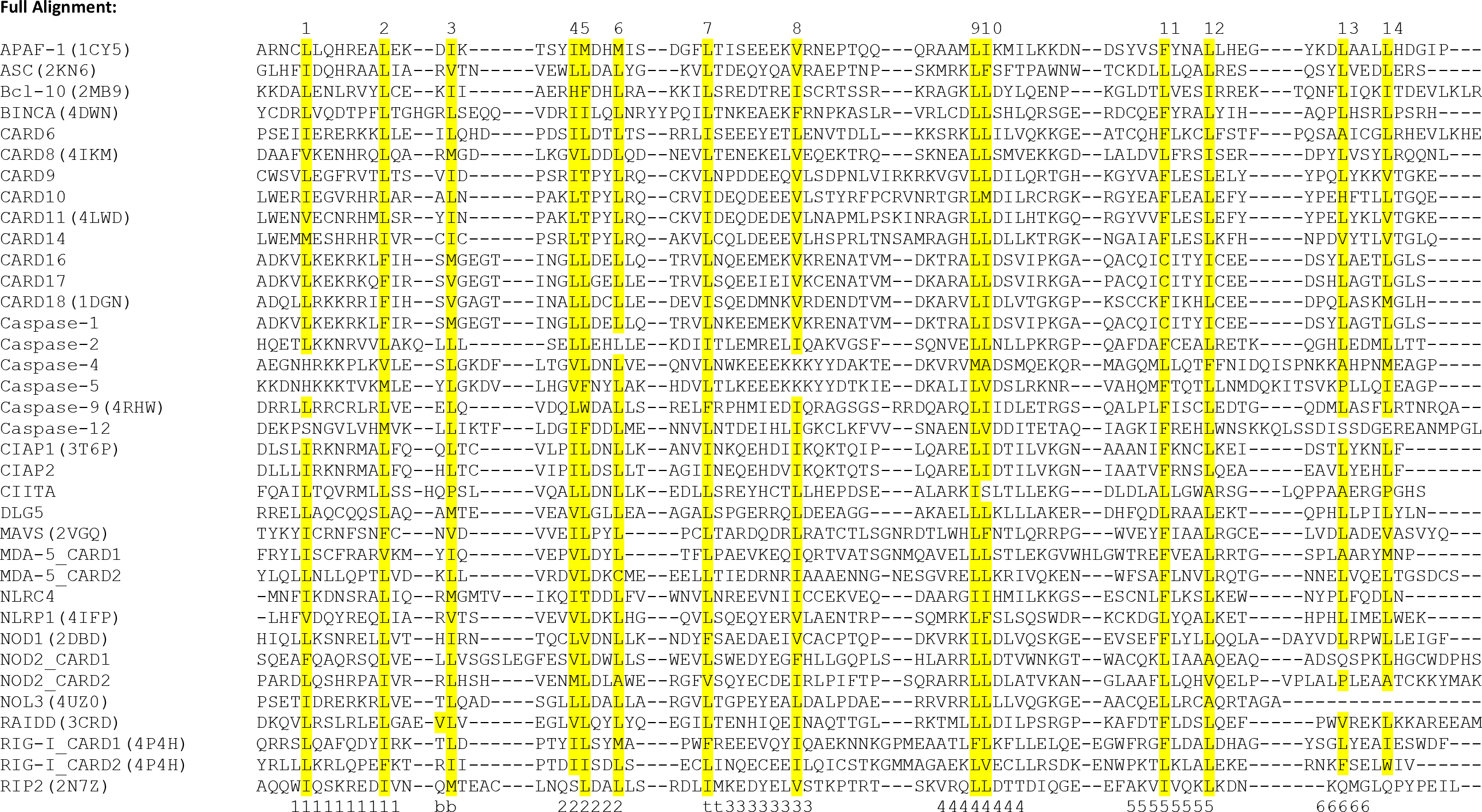
Alignment of 36 Human CARDs. Conserved hydrophobic residues, h1 – h14, are highlighted in yellow where known or predicted to be present in a sequence. Numbers underneath the alignment denote the positions of the six CARD helices. PDB codes used in the alignment for human CARDs are shown in brackets.

The majority of CARDs aligned well and could be fitted so that any insertions and deletions were placed outside of the likely helix positions while still fulfilling the requirement to create a hydrophobic core. Somewhat remarkably, the hydrophobic core identified in the preliminary alignment (Figure 1) showed a strong level of retention in all of the 36 CARD sequences contained in the final alignment (Figure 5). The final alignment is most reliable over the first five helices (marked with numbers 1-5 in Figure 5) and for the conserved hydrophobic residues. The size of the interhelical loops varies significantly between CARDs and is consequently more difficult to compare directly. The most likely hydrophobic residues for positions h13 and h14 are labelled in Helix 6 but structural information will be required to verify these in most cases. It has also been noted in previous studies (Hu et al., 2014) that the initiating methionine can contribute to the hydrophobic core, which suggests that the stability of recombinant constructs may benefit from beginning at the start of the protein sequence.

Analysis of the full CARD alignment identified a small number of notable deviations from the conservation of the hydrophobic core. These include: caspases-4 and −5 which both possess a lysine residue at position h8; Bcl-10 and RIPK2 which contain a histidine and a serine at position h4 respectively; CIITA, which contains a serine at position h10; and the presence of a threonine at h5 for NLRC4 and the highly similar group of CARDs formed from CARD9, 10, 11 and 14. We know that RIPK2 (PDB: 2NZ7), Bcl-10 (PDB:2MB9) and CARD11 (PDB: 4LWD) still adopt a CARD-like structure despite these substitutions indicating that the h4 and h5 positions have at least some flexibility in terms of the biochemical properties of the residue situated there. It remains to be seen whether caspase-4 and caspase-5 also adopt the classical CARD fold or whether the presence of a large positively charged side chain results in structural distortion. Of course, it may well be that such structural distortion is necessary for the CARDs of caspase-4 and caspase-5 to be able to act as intracellular sensors of LPS (Shi et al., 2014).

### Surface salt bridges are found in over half the CARDs

Eight of the sixteen structures analysed possess a salt bridge between an aspartate or glutamate on Helix 2 and a lysine or arginine in the 4-5 loop (Figure 6A, B). The alignment in Figure 5 shows that a further 12 CARDs contain appropriately charged residues in these positions, suggesting that 20 of the human CARDs maintain this salt bridge.

**Figure 6.**
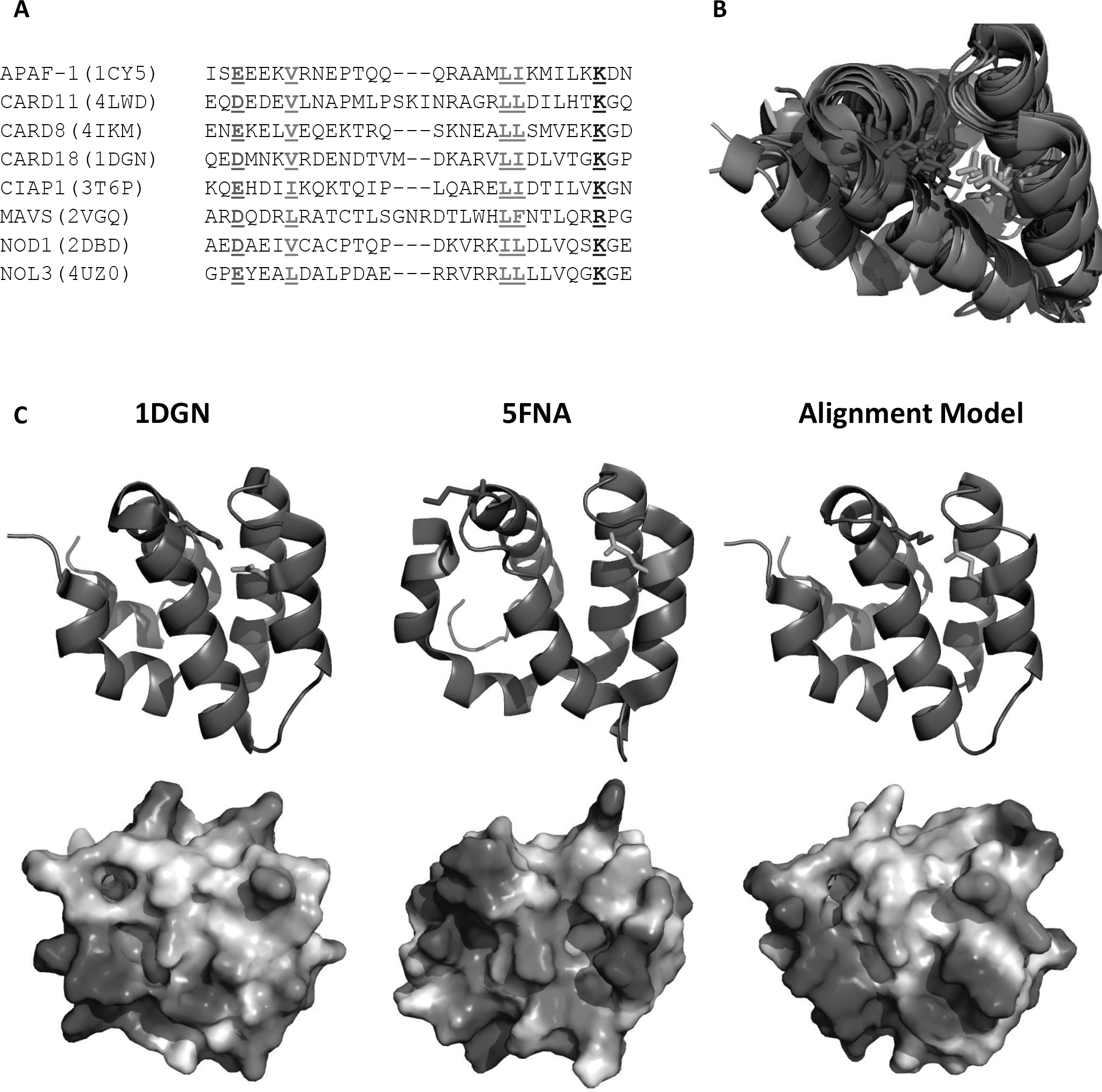
Many CARDs form a salt bridge between Helix 3 and the 4-5 Loop. (A) Eight of the currently available CARD structures contain a salt bridge between an acidic residue on Helix 3 (orange) and a basic residue in the 4-5 loop (blue). Conserved hydrophobic residues are shown in red and cover positions h8, h9 and h10. (B) Superposition of the eight CARD structures from the alignment in panel (A) with their salt bridges coloured. (C) Comparison of the positions of the CARD salt bridge from ICEBERG with the homology model of caspase-1 from 5FNA and a homology model based on our alignment.

It is plausible that these salt bridges may be important structurally and functionally. Certainly their potential formation needs to be considered when producing homology models and in the interpretation of site-directed mutagenesis data. The use of electron microscopy to study death domain filaments is becoming more common and in the absence of high-resolution monomeric structures models of these filaments are built using homology models of the individual domains. In a recent study (Lu et al., 2016) of the caspase-1 CARD filaments, homology models of the caspase-1 CARD were constructed based on the structure of the closely related CARD from CARD18 (ICEBERG; PDB 1DGN). These were then docked to an electron density map and provided a basis for understanding caspase-1 filament assembly. This study could arguably be further improved by the inclusion of a salt bridge in these homology models. Figure 6C shows the significant differences in residue position and surface electrostatic potential of caspase-1 homology models which either maintain or break this salt bridge, changes which may influence functional interpretation.

### Tyrosine phosphorylation in the type Ib interface

Both RIPK2 (Tigno-Aranjuez et al., 2010) and ASC (Hara et al., 2013) undergo tyrosine phosphorylation on a residue in their Type Ib patch – Y474 (RIPK2 human numbering) and Y144 (ASC murine numbering). These phosphorylation events are reportedly crucial for function, though they have not yet been observed in structural studies. Our alignment indicates that the CARDs of four other proteins, CARD6, NLRP1, NOL3 and NOD2 CARD1 have a tyrosine residue in the same position as ASC and RIPK2 (Figure 7A, B), suggesting that their activity might be modulated in a similar way and that post-translational modification may be a common way of mediating complex formation and downstream signalling for these proteins.

**Figure 7.**
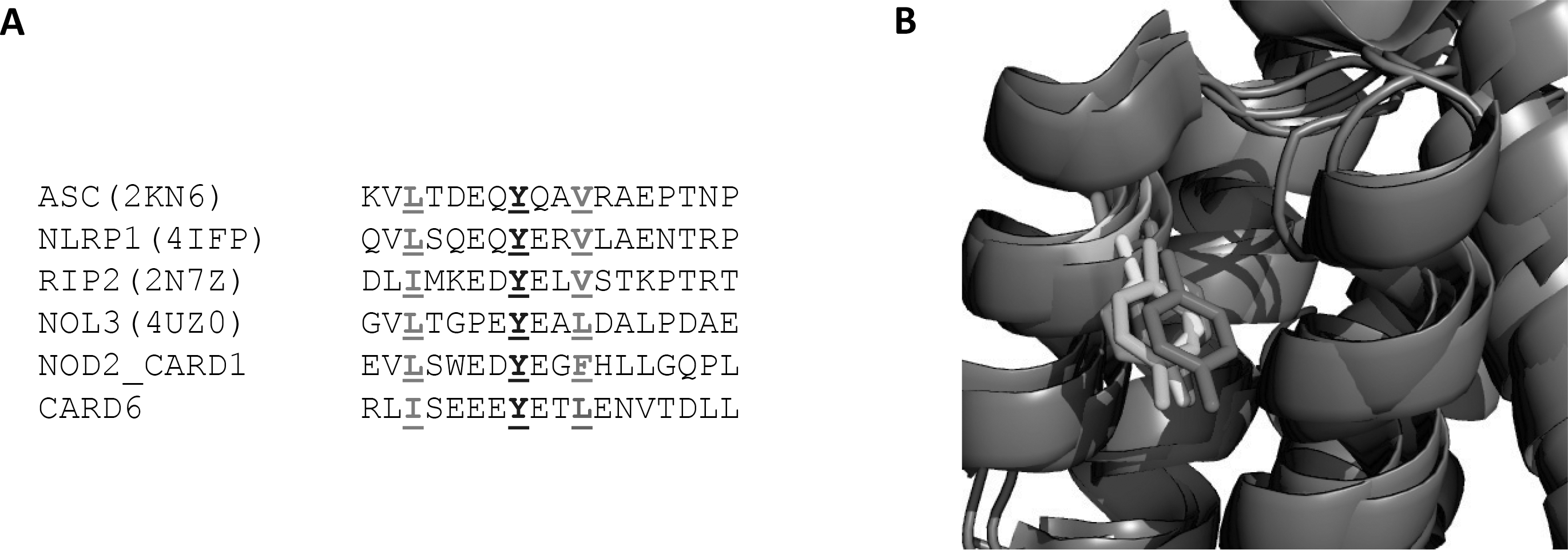
A tyrosine residue is found in the Type Ib patch of six human CARDs. (A) Alignment of six CARD sequences with the putatively phosphorylated tyrosine shown in blue. Conserved hydrophobic residues are shown in red and represent positions h7 and h8. (B) Overlay of four solved structures with tyrosines in the Type Ib patch.

The Type Ib patch in all six of these proteins also contains an aspartate residue and it may well be that tyrosine phosphorylation is necessary for the formation of a Type I interaction resembling that seen between caspase-9 and Apaf-1. In this instance positively charged and negatively charged surfaces interact around two key residues on each side (Qin et al., 1999). Further structural and functional studies of CARD:CARD interactions may need to use expression systems other than *E. coli* in order to investigate the importance of these phosphorylation events.

### The CARDs from human and murine NLRC3 and NLRC5 are pseudodomains

The structure of the N-terminus of murine NLRC5 has been solved by NMR. It forms an atypical CARD in that Helix 1a and Helix 6 are separated from the rest of the domain by extended loops and helix 3 is completely unstructured. Alignment of the human and murine NLRC5 sequences, based on the murine structure, shows that while most hydrophobic residues are conserved in both species, h1, h5, h8 and h9 are non-hydrophobic. The orthologous region of zebrafish NLRC5 contains the expected hydrophobic residues in these regions and in general aligned more readily with other CARDs (Figure 8A).

**Figure 8.**
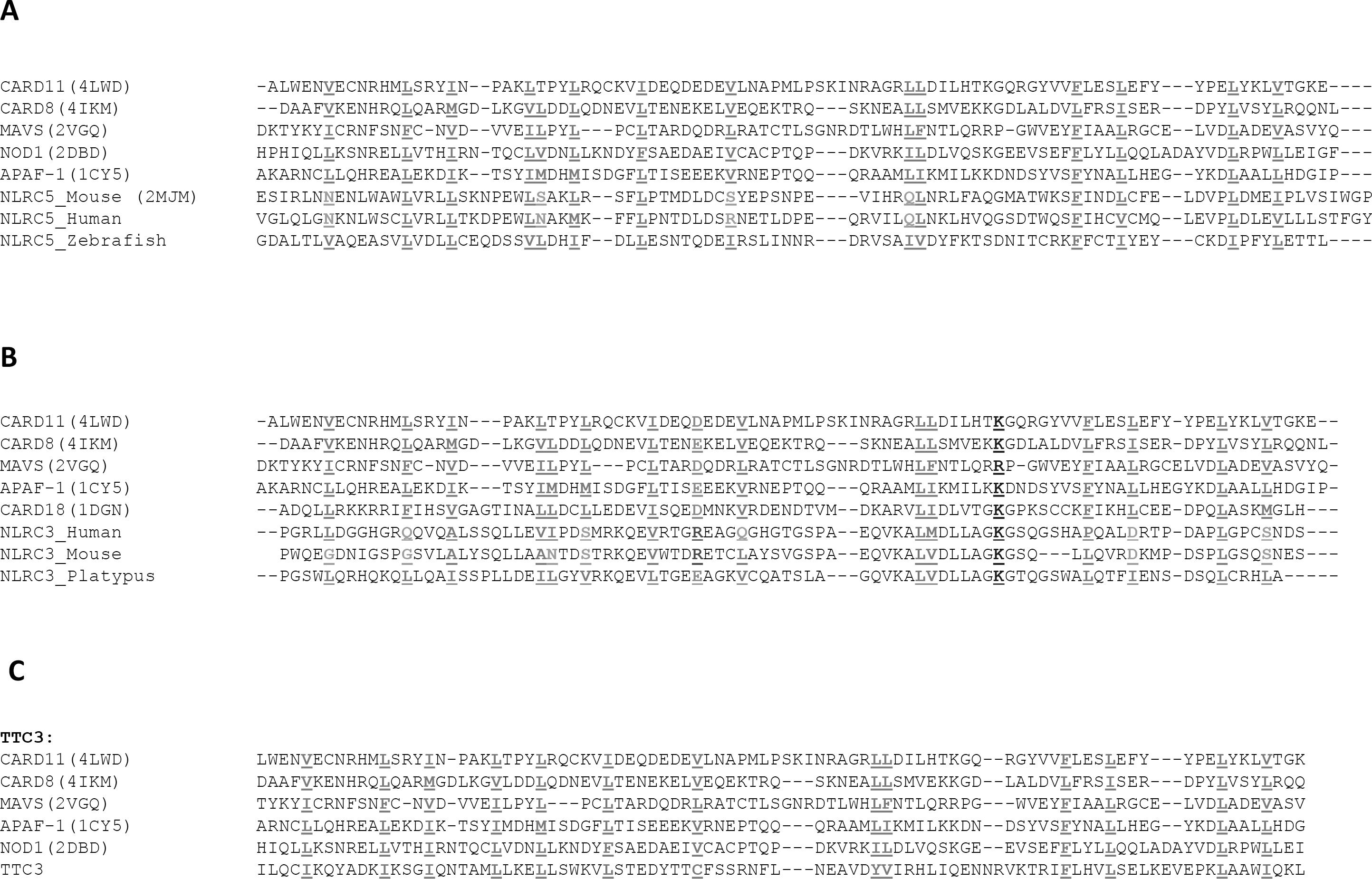
Human orthologues of NLRC5 and NLRC3 conform poorly to the conserved hydrophobic pattern. (A) Alignment of human, mouse and zebrafish NLRC5 orthologues with other CARDs. (B) Alignment of human, mouse and platypus NLRC3 orthologues with other CARDs. (C) The putatative CARD-containing protein TT3 shows conservation of the CARD hydrophobic core. In all panels residues contributing to the hydrophobic core are coloured red and presented in bold underlined text. Residues deviating from the hydrophobic core are light blue, bold and underlined. In panel B residues in the correct position to form a surface salt bridge are coloured orange (acidic), dark blue (basic) or purple (incompatible).

NLRC3 was examined in a similar manner. The zebrafish contains many NLRC3-like genes and so the NLRC3 orthologue from the platypus was used for comparison instead. Fold recognition analysis of the human NLRC3 sequence with FUGUE suggests that whilst many of the conserved hydrophobic residues are present in the human protein five – h2, h6, h8, h12, h14 – appear to have been altered (Figure 8B). The same pattern is seen for murine NLRC3 in which h1, h2, h5, h6, h12 and h14 are all substituted. In contrast, the platypus orthologue contains all of the expected hydrophobic residues and also appears to have the surface salt bridge commonly found in CARDs (Figure 8B) which is absent in both the murine and human NLRC3 CARDs.

The sequence deviation of the human and murine NLRC3 and NLRC5 CARDs, along with the nontypical structure of murine NLRC5 CARD indicate that they have become pseudodomains in humans and mice and have most likely lost some of their original functionality. Little is known about the roles of the NLRC3 and NLRC5 CARD, but our alignment shows apparent degeneration of these CARDs in humans and mice compared to their orthologues from certain vertebrates. The future study of CARD-containing proteins using other species might take these differences into account when drawing conclusions about protein function. It also suggests that too many substitutions in the hydrophobic core result in structural changes to the CARD fold. This has clear implications for downstream CARD function. This observation may have further implications for the CARDs of caspase-4 and caspase-5 which are substituted in the h1 and h8 positions and have uncertain helix 6 alignments. A disrupted or altered structure may help explain why these domains function as intracellular LPS receptors rather than the usual homotypic interaction motifs and structural scaffolds.

### The hydrophobic core can be used for identifying potential novel CARDs

Through use of PSI-BLAST and jackhmmer it is possible to find candidate CARDs in protein databases. However, the high sequence variability in the CARD superfamily can make these difficult to pick up on and confirm. One example from our search is Tetratricopeptide Repeat Domain 3 (TTC3), which came up as a low-confidence hit in multiple PSI-BLAST searches and was labelled as containing a possible CARD by FUGUE (Supplementary Table 1). The TTC3 sequence fits into the constraints of the conserved hydrophobic pattern (Figure 8C) and no other domains are predicted to overlap the CARD prediction, suggesting that it may indeed adopt a CARD-like fold, although of course ultimately structural confirmation will be required.

## Conclusion

We have used a combination of structure-based and sequence-derived information to create a global alignment of the human CARD superfamily. Almost all human CARDs were predicted to contain a set of fourteen hydrophobic residues and half of these also maintain a surface salt bridge. Deviations were most common in helix 6 while some CARDs such as that from RIPK2 also had single position exceptions to the hydrophobic pattern. The atypical structure of NLRC5, which is missing three conserved residues, suggests that too many differences can significantly alter the CARD fold, leading us to suggest that the CARDs of human and murine NLRC3 and NLRC5 are likely to be pseudodomains; whilst providing a potential explanation for the ability of the caspase-4 and −5 CARDs to bind LPS

Our investigation highlights the potential importance of hydrophobic residues, salt bridges and phosphorylation sites as a resource for successful design and interpretation of bioinformatic and experimental studies on the CARD superfamily. In particular, homology modelling and site-directed mutagenesis efforts may benefit from these observations. For example consideration of the potential structural roles of charged surface residues may aid the choice and design of appropriate expression systems to allow post-translation modifications and the production of functional, soluble and stable protein. It is highly likely that such an approach is also applicable to the other protein folds in the Death Domain superfamily.

## Materials and Methods

Fold identification and sequence alignments:

PSI-BLAST (http://blast.ncbi.nlm.nih.gov/Blast.cgi) (Altschul et al., 1997)and jackhmmer (https://www.ebi.ac.uk/Tools/hmmer/search/jackhmmer) (Finn et al., 2015) searches were performed using predicted CARD sequences from PFAM (Finn et al., 2016), extended by ten residues at both termini, as search terms (Supplementary Table 1). Each search was run to confluence with default settings. Fold recognition was performed using FUGUE (http://mizuguchilab.org/fugue/prfsearch.html) (Shi et al., 2001).

Automated sequence alignments of recovered CARD sequences were attempted using ClustalW2 (http://www.ebi.ac.uk/Tools/msa/clustalw2/) (Larkin et al., 2007), Clustal OMEGA (http://www.ebi.ac.uk/Tools/msa/clustalo/) (Sievers et al., 2011) and MUSCLE (http://www.ebi.ac.uk/Tools/msa/muscle/) (Edgar, 2004) using default settings.

Structural alignment was performed using, where available, one representative structure for each CARD (Supplementary Table 1). Generally, structures were chosen based on the best available resolution, though exceptions are detailed in the main text. Overlay and RMSD calculations were performed using either the “**super p1, p2**” command in PyMol (The PyMol Molecular Graphics System, Version 1.7.6.6 Schrödinger, LLC), where the first protein, p1, is superimposed onto the second protein, p2; or where the super command was not operational, “**super p1 & alt A+”, p2 & alt B+’’**” was used instead.

## Acknowledgements

This work was supported by the Wellcome Trust (WT085090MA), the Medical Research Council (U105960399) and a BBSRC Doctoral Training Grant.

## Supplementary Tables and Figures

**Supplementary Table 1 – Summary of Human CARDs used in this study.** Underlined proteins were found in the original Pfam database search. Proteins marked in italics are reported to possess either incomplete or irregular CARDs. PDB identifiers are only provided for human structures and those in bold are crystal structures, those in standard case are NMR structures. For structures that were part of a complex the relevant chain identifier is provided in parentheses.

**Supplementary Figure 1: Final alignment of the 17 CARD structures available from the PDB.** PDB identifiers for each structure are provided in brackets. Hydrophobic residues are highlighted yellow and numbered above the alignment. Helix position as per the Apaf-1 structure, PDB 1CY5, are marked by numbers below the alignment

